# Improved genome assemblies of plant-associated *Streptomyces* spp. as a resource for understanding plant pathogenicity in the genus

**DOI:** 10.64898/2026.08.18.745569

**Authors:** Brett A. Shelley, Matthew L. Fabian, Hien P. Nguyen, Alexandra J. Weisberg, Jeff H. Chang, Christopher R. Clarke

**Affiliations:** USDA Agricultural Research Service, Genetic Improvement for Fruits and Vegetables Lab, Beltsville, MD, USA; Oregon State University, Department of Botany and Plant Pathology, Corvallis, OR, USA

**Keywords:** *Streptomyces*, Thaxtomin, complete genome sequences, core genomes, plasmids, common scab

## Abstract

Common scab disease on potato is caused by members of more than 10 pathogenic *Streptomyces* species. Genome-enabled methods are being increasingly deployed to characterize *Streptomyces* that cause common scab disease of potato and other tuber and root crops. However, the study of phytopathogenic *Streptomyces* is constrained by the limited availability of high-quality genome sequences. Here we report improvements to the quality and completeness of genome assemblies for 12 pathogenic type strains of *Streptomyces* and six closely related non-pathogenic type strains. These assemblies have an average N50 of 7.4 Mbp and with BUSCO scores all greater than 98.5%. Analyses showed that the genomes of phytopathogenic *Streptomyces* are consistently among the largest *Streptomyces* genomes sequenced and, relative to those of non-pathogenic strains, are more enriched in genes involved in carbohydrate and amino acid metabolism. Plasmids were not consistently detected across assemblies, suggesting that they are not conserved across species and are not necessary for pathogenicity. Furthermore, comparisons of genome assemblies among both closely and distantly related strains revealed multiple rearrangements within linear chromosomes and reduced synteny near telomeric regions. These improved genome assemblies, many of which correspond to type strains, provide valuable resources for advancing our understanding of the pathogenicity in the genus.

## Introduction

The Gram-positive *Streptomyces* genus consists of over 900 species with standing in nomenclature (LPSN.dsmz.de, [1], encompassing a wide ecological range from saprophytes to plant pathogens. Among phytopathogenic species, the production of the phytotoxin Thaxtomin A is strongly associated with the ability to cause potato common scab disease, one of the most economically important diseases affecting potato crops [2–9]. Although Thaxtomin A is central to common scab disease, several species of *Streptomyces* have been reported to cause other scab diseases of potato, such as netted scab, without producing Thaxtomin A. These other scab diseases are generally considered less severe than common scab [10–14]. Phytopathogenic *Streptomyces* that cause potato common scab and other scab diseases of potato exhibit tremendous phylogenetic diversity [2,15–17]. Recent studies have also identified several novel species of *Streptomyces* phytopathogens, revealing even more diversity than previously recognized [18–21].

Complete genome sequences are critical resources for gaining a fundamental understanding of the genetic basis of virulence and diversity among phytopathogenic *Streptomyces*, and for developing effective disease management strategies*. Streptomyces* strains, including phytopathogenic strains, have a single linear chromosome and sometimes one or more plasmids [22,23] [24]. While plasmids are known to contribute to disease in other Gram-positive bacteria plant pathogens [17,25,26], presence and conservation of plasmids among potato common scab *Streptomyces* pathogens is unknown. There are numerous publicly available genome sequences of non-pathogenic *Streptomyces*, several of which are complete, but there is a paucity of complete or near-complete genome assemblies for phytopathogenic *Streptomyces* (**Figure 1A**). Queries of *Streptomyces* genome sequences from online repositories yield a wide range in the quality of genome assemblies, with most being of low quality [27,28]. Some assemblies have more than 1,000 scaffolds, often with no additional work that advances the quality or completeness of the assembly. Additionally, other entries contain complete or near-complete genome sequences but correspond to a strain that is not the type-strain of the species. For example, the two highest quality available genome sequences among phytopathogenic strains of *Streptomyces* are *S. scabiei* strain 87.22 and *S. acidiscabies* strain NRRL B-16521, neither of which are species type-strains. Per NCBI, among the genome sequences of the 14 reported pathogenic species and 6 closely related non-pathogenic strains, only 2 type-strain genome assemblies are nearly complete with fewer than 10 contigs: *S. griseiscabiei* NRRL B-2795 [19] and *S. puniciscabiei* NRRL B-24456.

**Figure 1.**
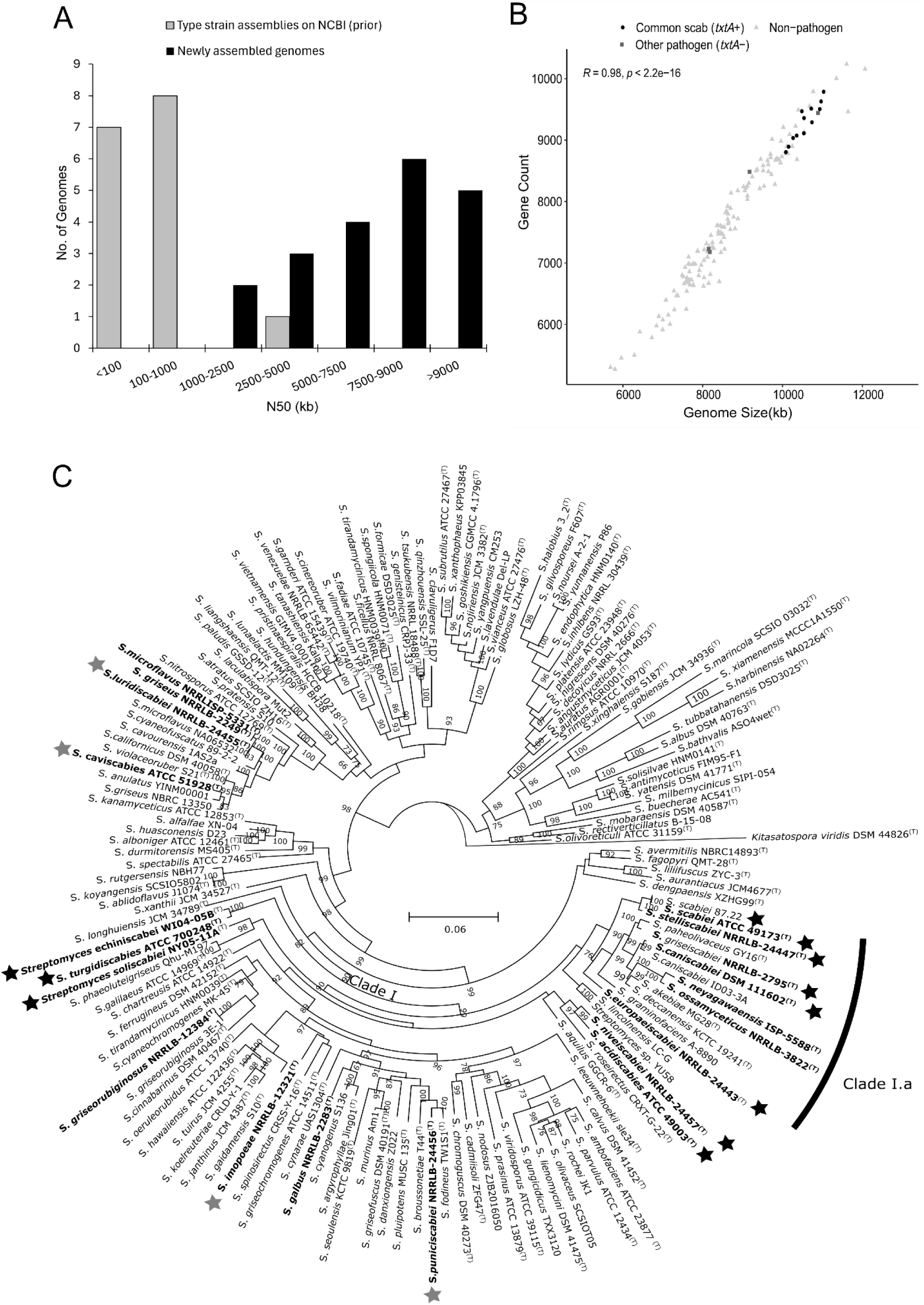
Phylogenetic relationship and genome status of complete and near-complete *Streptomyces* genomes. **(A)** Distribution of the genome assembly N50 for *Streptomyces* type-strains sequenced in this work before and after re-sequencing, including *S. echiniscabiei* WI04-05B *and S. soliscabiei* NY05-11A. Black bars represent those on NCBI and grey bars represent the improved assemblies from this study. **(B)** Comparison of Gene Counts to Genome Size (bp) of complete or near-complete, annotated genomes of *Streptomyces* using Prokka. *Streptomyces* were grouped as non-phytopathogens (NP), common scab pathogens (*txtA*+), or other pathogens (lacking *txtA* and reported to cause diseases other than common scab). **(C)** Maximum likelihood phylogeny based on five concatenated gene sequences (*atpD, gyrB, recA, rpoA, trpD*) of the 153 Complete Genome Sequences used in this analysis. Tree was generated using default parameters of iqtree2 within automlsa2. Pathogens are indicated with stars, *txtA-*positive strains are indicated by a black star. Strains sequenced in this work or [21] are in bold font. Tree was rooted to the outgroup *Kitastospora viridis* DSM44826^T^. Strains missing more than three of the marker genes were excluded from the analysis.

There are several challenges with generating high quality *Streptomyces* genome assemblies. Some common features of *Streptomyces* genomes pose challenges for generating complete, high quality genome assemblies, include: an unusually rich G+C content (>70% mol); a typically large genome size of 8 Mbp; and the presence of several highly repetitive regions [28]. High G+C content can also compromise assembly accuracy, as well as introduce bias to PCR amplification, while large and repeat-rich genome sequences are known to contribute to fragmentation of assemblies. Moreover, these challenges are amplified when using short-read Illumina sequencing, which was the technology of choice for most current genome assemblies of *Streptomyces* [27,29]. Long read, polymerase-free sequencing with PacBio CLR or Oxford Nanopore (ONT) technologies can help overcome these challenges, however long read sequencing can introduce higher error rates for base call accuracy. An alternative approach is the hybrid assembly, which integrates both short and long read sequences. Hybrid assemblies have the potential to correctly resolve complex regions into a complete assembly while maintaining high per-base sequence accuracy [29].

Improved genome assemblies for phytopathogenic *Streptomyces* are desirable for advancing our understanding of the basis and diversity of phytopathogenicity within the genus. Given the large number of often distantly related phytopathogenic *Streptomyces*, numerous high quality reference genome assemblies are needed to cover the diversity of known pathogenic lineages. Here we report complete or nearly complete genome assemblies for 12 pathogenic *Streptomyces* type strains and 6 additional non-pathogenic type strains in closely related or sister taxa.

## Methods

### DNA Extraction

*Streptomyces* cultures were maintained on Arginine Glycerol Agar Sucrose (AGS, 1 g arginine, 12.5 g glycerol, 1 g K_2_HPO_4_, 1 g NaCl, 0.5 g CaCO_3_, 0.5 g MgSO_4_*7H_2_0, and 1 mL micronutrients solution L^-1^ ddH_2_O, pH 7) for 14 days. A total of 4 mL sterile water was used to remove bacteria from the plate, and the suspension was used to inoculate 30 mL of Yeast Malt Extract (YME; 4g yeast extract, 10g malt extract, and 4 g dextrose L^-1^ ddH_2_O). After 48 h of growth at 28 °C with shaking at 200 rpm, cultures were pelleted, and genomic DNA was extracted using the HMW Promega DNA wizard kit, following the protocol for Gram-positive bacteria (Promega, CA, USA) with modifications previously described [2].

### Generation of Sequencing Reads

For Illumina sequencing, DNA libraries were prepared by the Center for Quantitative Life Sciences (CQLS) at Oregon State University or by Genewiz/Azenta (NJ, USA) and sequenced as 150 bp paired-end reads on Illumina HiSeq 3000 sequencers. For PacBio sequencing, libraries were prepared by Genewiz/Azenta and sequenced on the Sequel platform. Additional long read sequencing was completed using a MinION Mk1C device with an R.9.4.1 flow cell (Oxford Nanopore, Cambridge, UK). Genomic libraries for ONT sequencing were constructed using the ONT rapid barcoding kit (SQK-RBK004) and/or ligation sequencing kits (SQK-LSK109/110). Publicly available sequences within the Short Read Archive (SRA) were downloaded from NCBI [30].

### Genome Assembly

Adapters and barcodes were trimmed using Porechop v. 0.2.4 and the reads were combined into single FASTQ files. Long-read sequences were assembled using Flye v. 2.9 with the default options and “--genome-size 10m” [31]. When both Oxford Nanopore reads and PacBio reads were available for the same genome, they were combined and assembled using Flye with the following modifications: “--pacbio-raw $PBREADS $ONTREADS --iterations 0”. Flye was then re-run, with the following modifications: “--pacbio-raw $PBREADS --resume-from polishing --genome-size 10m”. The long read assembly was then merged with fastp - corrected Illumina reads via Unicycler v. 0.5.0 with SPAdes v. 3.15 with the following modifications: “– existing_long_read_assembly $FlyeAssembly --linear_seqs 1” [22,28,29,32,33], as well as “–mode bold”. Contigs that contained the PhiX gene were manually removed from the genome assemblies. The completeness of the genome assemblies was determined by using Benchmarking Universal Sing-Copy Orthologs (BUSCO) v. 5.2.2 [34] with the lineage dataset streptomycetales_odb10. The hybrid genome assembly pipelines are shown in **Figure S1**.

### Annotation and pangenome identification

All complete *Streptomyces* reference genome sequences (n=133) were downloaded from NCBI on February 6^th^, 2024. All genome sequences for *S. scabiei, S. caniscabiei, S. acidiscabies, and S. stelliscabiei* were downloaded from NCBI on May 17^th^, 2024. The newly generated genome assemblies were annotated using Prokka v. 1.14.6 [35] with default parameters, and genome assemblies downloaded from NCBI were reannotated in the same fashion. The pangenome from *Streptomyces* complete or nearly complete genome assemblies was generated using PIRATE [36] using the following modifications: “ -t 256 -a -r -k --cd-mem 100000".

### Genome-wide association study (GWAS) for common scab pathogenicity

All genome assemblies were categorized based on the reported pathogenicity of corresponding strains and presence/absence variation of *txtA,* used as a marker for the Thaxtomin A gene cluster. Strains were sorted into three groups: 1) *txtA*-positive common scab pathogens, 2) *txtA-*negative “other” scab pathogens, and 3) non-pathogens (*txtA*-negative). Annotated gene sequences for txtA-positive (n=12) and non-pathogenic (n=137) strains were provided as input to OrthoFinder [37], which, using default options, generated hierarchical orthogroups (HOgs) and a rooted, species-level Newick tree. The complete set of HOgs (n = 52,660) and the Newick tree were used in a GWAS for presence of the *txtA* gene using Scoary2 [38] with default options. HOgs with a Bonferroni-adjusted p-value (Fisher’s exact post hoc test) < 0.05 were identified as significantly associated with common scab pathogenicity. Significant associations were further identified on the basis of pairwise comparison testing and post hoc binomial test p-values < 0.05 [39]. Orthogroups with significant associations with pathogenicity were retrieved, along with their associated amino acid sequences. Representative sequences for each HOg were assigned protein accession numbers from the RefSeq Select database via “blastp” [40] with the following parameters: “-evalue 1e-5 -outfmt 6 -max_target_seqs 1 -taxids 1883”. Gene names were assigned to protein accession numbers via edirect [41]. For enrichment analysis of HOgs that have a significant positive association with common scab pathogenicity, representative amino acid sequences for each of the underlying orthogroups for all HOgs were submitted to BlastKOALA [42] for assignment of KEGG Orthology identifiers from the “Prokaryotes” database. Orthogroup counts were summarized at the KEGG subcategory and pathway levels, and enrichment for each level was expressed as (orthogroup count for significant HOgs) * (background orthogroup count)^-1^.

### Plasmid analysis

Assembly graphs were examined via Bandage [43], and contigs identified by Unicycler as circular (i.e., “circular=TRUE”) were identified as candidate plasmids. Additionally, linear contigs with coverage depth ≥ 2.00 were characterized as candidate plasmids. Assembled genome sequences were analyzed using plasmidFinder [44] with default parameters for Gram-positive bacteria. All putative plasmids were analyzed via BLAST in comparison to previously reported *Streptomyces* plasmid sequences. Sequences of putative plasmids were then mapped against the genome assembly sequence of *Streptomyces scabiei* 87.22 (GCF_000091305.1) and queried for the presence of sequences of *txtA* and *txtB,* and core housekeeping genes 16s rRNA, *gyrB*, and *trpB*. Contigs that lacked homology to these sequences and did not map to the 87.22 genome assembly were designated as putative plasmids. Putative plasmids were also examined for regions of homology to the *repA* and *repB* sequences from the *S. violaceoruber* pSV1 plasmid (JQ659191.1).

### Synteny mapping

Genome assemblies were aligned to the *Streptomyces scabiei* 87.22 complete assembly via minimap2 with the parameter “-ax asm20” [45]. Where necessary, and after accounting for alignment orientation and contig breaks, contigs were concatenated into a single chromosomal contig. Small contigs for which alignment orientation and order could not be resolved were excluded from concatenation. The single-chromosome assemblies were aligned using the “nucmer” utility within MUMmer [46], and with the “--maxmatch” parameter. Alignments were filtered using the “delta-filter” utility and the parameters “-m -i 60 -l 100”. Alignment coordinates were generated using the “show-coords” utility and the parameter: “-THrd”. Synteny mapping files were generated using SyRI [46] with default settings. Synteny plots were generated using plotsr [47] with default settings.

The complete genome assemblies were aligned using NUCmer embedded in the CSAR (contig scaffolding using algebraic rearrangements) webserver (http://lu168.cs.nthu.edu.tw/CSAR-web/index.php) [48]. For this analysis, putative plasmid contigs were removed and only the predicted chromosomal contigs were used. The location of TR1 was aligned to each of the newly assembled genome assemblies with progressiveMauve [49], using the default settings with *S. scabiei* strain 87.22 as a reference. When inversions were observed across an entire contig in the final alignment, the contig was reverse complemented, re-ordered and re-run.

## Results

*Improved genome assemblies of phytopathogenic and closely related* Streptomyces *type strains*

We used short and long read sequences and existing sequence read archive (SRA) data to produce complete or nearly complete genome assemblies for 18 selected strains via an optimized hybrid assembly pipeline (**Figure S1, Table 1**). Two additional genome assemblies of recently described novel species (*S. soliscabiei* NY05-11A and *S. echiniscabiei* WI04-05B) of phytopathogenic *Streptomyces* sequenced in parallel and previously reported [21] were included in downstream analyses. For several of the genome assemblies, inclusion of publicly available SRA data degraded the quality of the assembly and therefore were not used. The average size of the genomes of the newly sequenced phytopathogenic strains was 10.1 Mbp, with an average N50 of 7.4 Mbp, significantly improving the assembly quality of all genomes for which publicly available assemblies were previously available (**Figure 1A**). BUSCO (Benchmarking Universal Single-Copy Orthologue) analysis of the re-sequenced and assembled genomes predicted high levels of completeness, with all scores greater than 98.5%. Final assembly statistics are reported in **Table 2** and **Table S1**. Quality and completeness of our assemblies were compared to publicly available assemblies within the same species group. The number of missing and fragmented BUSCOs per genome sequence were plotted against the assembly N50 values for all genomes of the four pathogenic species groups (*S. scabiei, S. acidiscabies, S. caniscabiei,* and *S. stelliscabiei*) that contained more than 10 publicly available genome assemblies on NCBI. In general, there was a negative correlation, with N50 inversely related to the number of missing or fragmented BUSCOs (**Figure S2**). Based on these metrics, the newly generated assemblies of the type strains are an improvement on other available genome assemblies for each of the species groups. Among the newly assembled and publicly available, complete/near-complete genomes of *Streptomyces,* there is a strong correlation (*R=*0.98, *p <* 2.2e^-16^) between gene count (coding sequences (CDS), rRNA, tRNA) and genome size, as expected (**Figure 1B**). Interestingly, the *txtA-*positive pathogens (*i.e.*, common scab pathogens) were among the largest genomes (>10 Mbp) across all *Streptomyces*. There was a broad distribution of genome size across the other *Streptomyces* genomes ranging from < 6 Mbp to >12 Mbp. To determine the phylogenetic context of the strains with complete and nearly complete genome assemblies used in this work, we generated a maximum likelihood tree using established multi-locus sequence analysis (MLSA) loci (**Figure 1C**). The *txtA-*positive (common scab) pathogens are distributed into three main clades with several closely related non-pathogenic *Streptomyces* within the main *S. scabiei* clade, as expected [2].

**Table 1.** Assembly Methods and Read Inputs for assembled genomes.

| Species | Isolate | Genbank Accession | Assembly Method | Long Read Types<sup>1</sup> | Illumina Read Source | Coverage (X) |  |  |
| --- | --- | --- | --- | --- | --- | --- | --- | --- |
|  |  |  |  |  |  | PacBio | ONT | Illumina |
| *S. scabiei* | ATCC49173 | JBPJFT000000000 | Flye+Unicycler | ONT, PB | This work, [50] | 107 | 15 | 46 |
| *S. europaeiscabiei* | NRRL B-24443 | JBPJFK000000000 | Flye+Unicycler | ONT, PB | This work, [50] | 60 | 135 | 67 |
| *S. stelliscabiei* | NRRL B-24447 | JBPJFJ000000000 | Flye+Unicycler | ONT, PB<sup>2</sup> | This work, [50] | 291 | 66 | 71 |
| *S. griseiscabiei* | NRRL B-2795 | JBPJFN000000000 | Flye+Unicycler | ONT | [50] | n/a | 67 | 103 |
| *S. caniscabiei* | NE06-02D | JBPJFP000000000 | Flye+Unicycler | ONT | [9] | n/a | 38 | 48 |
| *S. turgidiscabies* | ATCC700248 | JBPJFR000000000 | Flye+Unicycler | ONT, PB | This work, [50] | 131 | 39 | 37 |
| *S. acidiscabies* | ATCC49003 | JBPJFU000000000 | Flye+Unicycler | ONT, PB | This work, [50] | 137 | 17 | 20 |
| *S. niveiscabiei* | NRRL B-24457 | JBPJFH000000000 | Flye+Unicycler | ONT, PB | This work | 109 | 50 | 65 |
| *S. luridiscabiei* | NRRL B-24455 | JBPZSP000000000 | Flye+Unicycler | ONT, PB | This work | 113 | 8 | 67 |
| *S. puniscabiei* | NRRL B-24456 | JBPJFI000000000 | Flye+Unicycler | ONT, PB<sup>3</sup> | This work | 567 | 296 | 52 |
| *S. caviscabies* | ATCC51928 | JBPJFS000000000 | Flye+Unicycler | ONT | [50] | n/a | 49 | 75 |
| *S. ipomoeae* | NRRL B-12321 | CP182305 | Flye+Unicycler | PB | This work | 113 | n/a | 82 |
| *S. griseorubiginosus* | NRRL B-12384 | JBPJFL000000000 | Flye+Unicycler | ONT | [50] | n/a | 10 | 65 |
| *S. griseus subsp. alpha\** | NRRL B-2249 | JBPZSQ010000000 | Flye+Unicycler | ONT, PB | [50] | 232 | 27 | 174 |
| *S. microflavus* | NRRL ISP-5331 | JBPZSQ000000000 | Flye+Unicycler | ONT | This work | n/a | 23 | 26 |
| *S. neyagawaensis* | NRRL ISP-5588 | JBPJFQ000000000 | Flye+Unicycler | ONT | This work | n/a | 75 | 24 |
| *S. galbus* | NRRL B-2283 | JBPJFO000000000 | Flye+Unicycler | ONT | This work, <sup>4</sup> | n/a | 37 | 50 |
| ***S. ossamyceticus*** | **NRRL B-3822** | JBPJFM000000000 | Flye+Unicycler | ONT | This work | n/a | 385 | 29 |

<sup>1</sup>PB, PacBio; ONT; Oxford Nanopore Technologies; Unless indicated, the source of the long read sequence reads is this work.
<sup>2</sup> SRR9008389, SRR9008390
<sup>3</sup> SRR12604680
<sup>4</sup> SRR7783799
\*Reassigned to *S. microflavus*

**Table 2.** Statistics for 18 newly generated *Streptomyces* complete or near-complete genome assemblies.

| Species | Isolate | Previous Assembly Statistics |  |  |  |  |  |  | New Assembly Statistics |  |  |  |  |  |
| --- | --- | --- | --- | --- | --- | --- | --- | --- | --- | --- | --- | --- | --- | --- |
|  |  | Txt-AB | Contigs | Assembled Genome Size (Mbp) | Contigs N50 (kbp) | GC Content (%) | No. CDS | BUSCO completeness (%) | Contigs | Assembled Genome Size (bp) | Contigs N50 (bp) | GC Content (%) | No. CDS | BUSCO completeness (%) |
| *S. scabiei* | ATCC49173 | Yes | 120 | 10.0 | 205.6 | 71.5 | 8639 | 99.7 | 3 | 10,081,288 | 9,936,670 | 71.5 | 8688 | 99.7 |
| *S. europaeiscabiei* | NRRL B-24443 | Yes | 1056 | 10.0 | 16.4 | 70.5 | 8789 | 96.8 | 4 | 10,531,514 | 4,162,400 | 70.6 | 8972 | 99.7 |
| *S. stelliscabiei* | NRRL B-24447 | Yes | 991 | 10.0 | 19.0 | 71.0 | 8845 | 94.5 | 4 | 10,351,907 | 6,791,917 | 71.1 | 8965 | 99.7 |
| *S. griseiscabiei* | NRRL B-2795 | Yes | 8 | 10.6 | 4000 | 71.5 | 9390 | 98.5 | 5 | 10,930,739 | 8,469,695 | 71.4 | 9391 | 98.5 |
| *S. caniscabiei* | NE06-02D | Yes | 84 | 10.9 | 446 | 71.5 | 9516 | 99.8 | 3 | 10,963,976 | 10,788,157 | 71.2 | 9523 | 99.7 |
| *S. turgidiscabies* | ATCC700248 | Yes | 71 | 10.7 | 289.6 | 70.0 | 9385 | 99.7 | 4 | 10,717,662 | 10,438,753 | 69.9 | 9409 | 99.7 |
| *S. acidiscabies* | ATCC49003 | Yes | 132 | 10.9 | 192.6 | 70.5 | 9575 | 99.1 | 5 | 11,027,707 | 7,538,445 | 70.6 | 9680 | 99.1 |
| *S. niveiscabiei* | NRRL B-24457 | Yes | 1568 | 9.5 | 10.1 | 71 | 8771 | 91.8 | 8 | 10,536,631 | 2,893,404 | 71.3 | 9259 | 98.9 |
| *S. imopoeae* | NRRL B-12321 | No | 213 | 10.9 | 174 | 70.5 | 9371 | 99.5 | 1 | 10,886,267 | 10,886,267 | 70.3 | 9336 | 99.5 |
| *S. luridiscabiei* | NRRL B-24455 | No | 821 | 7.9 | 18 | 71.5 | 7044 | 96.3 | 4 | 8,177,249 | 5,135,925 | 71.4 | 7080 | 99.5 |
| *S. puniscabiei* | NRRL B-24456 | No | 688 | 8.8 | 25.4 | 71.5 | 8144 | 94.7 | 3 | 9,171,190 | 8,082,017 | 71.4 | 8368 | 99.5 |
| *S. caviscabies* | ATCC51928* | No |  |  |  |  |  |  | 2 | 8,146,051 | 8,122,383 | 71.8 | 7121 | 98.7 |
| *S. griseorubiginosus* | NRRL B-12384* | No |  |  |  |  |  |  | 6 | 9,821,049 | 5,017,732 | 70.9 | 8573 | 99.8 |
| *S. griseus* subsp. *alpha* | NRRL B-2249* | No |  |  |  |  |  |  | 2 | 8,623,867 | 8,384,118 | 71.2 | 7604 | 99.5 |
| *S. microflavus* | NRRL ISP-5331* | No |  |  |  |  |  |  | 8 | 8,442,654 | 4,292,933 | 71.2 | 7497 | 98.5 |
| *S. neyagawaensis* | ISP-5588* | No |  |  |  |  |  |  | 9 | 10,082,296 | 2,118,439 | 71.4 | 8677 | 99.8 |
| *S. galbus* | NRRL B-2283* | No |  |  |  |  |  |  | 12 | 7,874,305 | 4,730,767 | 73 | 7094 | 98.3 |
| *S. ossamyceticus* | NRRL B-3822 | No | 1107 | 9.3 | 16.3 | 71 | 8224 | 93.4 | 7 | 10,080,667 | 2,482,103 | 71.3 | 8661 | 99.3 |

\*First reported assembly of strain.

### Multiple pathways are enriched in phytopathogenic Streptomyces

To identify gene content characteristics across the *Streptomyces* genus, 133 complete, annotated genome assemblies for *Streptomyces* were downloaded from NCBI and analyzed alongside the 20 newly assembled genomes, representing 149 distinct species (**Table S2**). A total of 98,172 orthologous gene families containing 714,924 gene sequences were identified across the entire set of 153 complete genome assemblies of *Streptomyces* (**Table S3**). A plot of ortholog conservation among strains displays a typical pattern, with the majority of orthologs present in only one or a few of the analyzed genome sequences (**Figure S3A**). Most *Streptomyces* genome assemblies have fewer than 7,500 predicted members of orthologous gene clusters; however, *Streptomyces* species harboring Thaxtomin A biosynthetic genes and closely related strains consistently have genomes exceeding 7,500 total predicted genes (**Figure S3B)**, consistent with their larger genome sizes.

A genome-wide association study (GWAS) was performed using *txtA* presence/absence as the trait of interest, comparing *txtA*-positive pathogenic strains to non-pathogenic strains, in order to identify genes potentially associated with phytopathogenicity. Because there is ambiguity around the disease symptoms of the *txtA-*negative strains that are reported to cause netted scab and other tuber diseases, we excluded these genome sequences from these analyses. In total, 546 hierarchical orthogroups (HOgs), including the Thaxtomin A biosynthetic genes, were identified as significantly enriched among the genome assemblies of the *txtA*+ pathogenic *Streptomyces* (**Table S4**). All HOgs were assigned KEGG orthology identifiers to discover functional commonalities among HOgs enriched among *txtA*-positive strains. As conveyed by both total gene count and relative enrichment among pathogen-associated HOgs, the KEGG subcategories most associated with phytopathogenicity were xenobiotics metabolism, carbohydrate metabolism, amino acid metabolism, secondary metabolites, and membrane transport (**Figure 2A**). A total of 66 associated KEGG pathways were identified (**Table S5**), and the top 30 enriched KEGG pathways were analyzed for relative enrichment (**Figure 2B**). Within the amino acid metabolism KEGG subcategory, highly enriched KEGG pathways corresponding to glycine, serine, threonine, arginine, tyrosine, and lysine metabolism were identified. Additionally, underpinning the carbohydrate metabolism KEGG subcategory were multiple KEGG pathways corresponding to starch and simple sugars, in addition to the “glycan degradation” pathway. While some enriched pathways (e.g., “bacterial chemotaxis”) do not make sense in the context of *Streptomyces*, they were observed to be supported by a very small number of genes.

**Figure 2.**
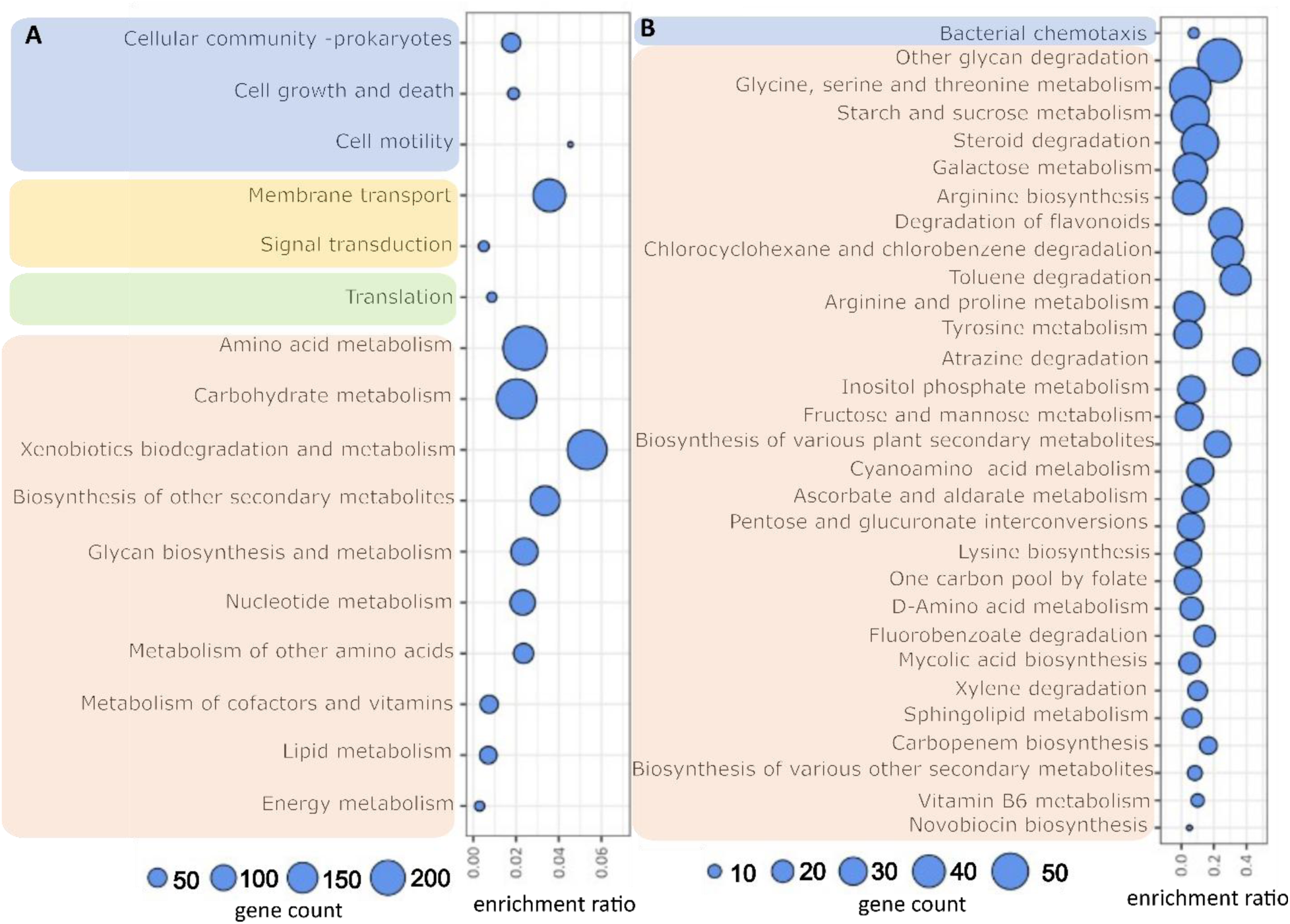
KEGG pathways assigned to genes in *txtAB*+ strains significantly associated with common scab. **(A)** KEGG pathway subcategory enrichments. **(B)** The top thirty enriched KEGG pathways. Gene orthologous groups (Ogs) identified via OrthoFinder were assigned to KEGG Orthology identifiers via BlastKOALA and summarized by KEGG pathway. “Enrichment ratio”, the ratio of the number of common scab-associated hierarchical Ogs (HOgs) to total Ogs for the corresponding KEGG subcategories or pathways. Shading corresponds to KEGG pathway categories: blue, “Cellular Processes”; yellow, “Environmental Information Processing”; green, “Genetic Information Processing”; and red, “Metabolism.” “Gene count”, the number of underlying genes, in common scab-causing strains, for the corresponding HOgs.

### Identification of putative plasmids in phytopathogenic Streptomyces

Plasmids have been reported in several species of *Streptomyces* and contribute to disease in other Gram-positive bacteria plant pathogens [25,26]. To assess whether plasmids play an important role in common scab, genome sequences were analyzed using PlasmidFinder, however no plasmids were predicted when the Gram-positive filter was utilized. Alternatively, plasmids were predicted by identifying circular contigs within the Bandage maps (**Figure S4**), and a total of nine predicted plasmids were identified among six common scab-causing strains (**Table 3**). Similarly, several linear contigs were identified that yielded sequencing depths greater than 2.00 (**Table 3**), potentially indicative of linear plasmids. However, only a few of the putative circular or linear plasmids also contained genetic loci with similarity to known plasmids. Of the putative plasmids identified, none contained homologous regions for *repA* and *repB* sequences. There was not a single conserved putative plasmid among the genome assemblies of the phytopathogenic *Streptomyces*, suggesting that plasmids are not necessary for virulence of common scab-causing *Streptomyces*.

**Table 3.**
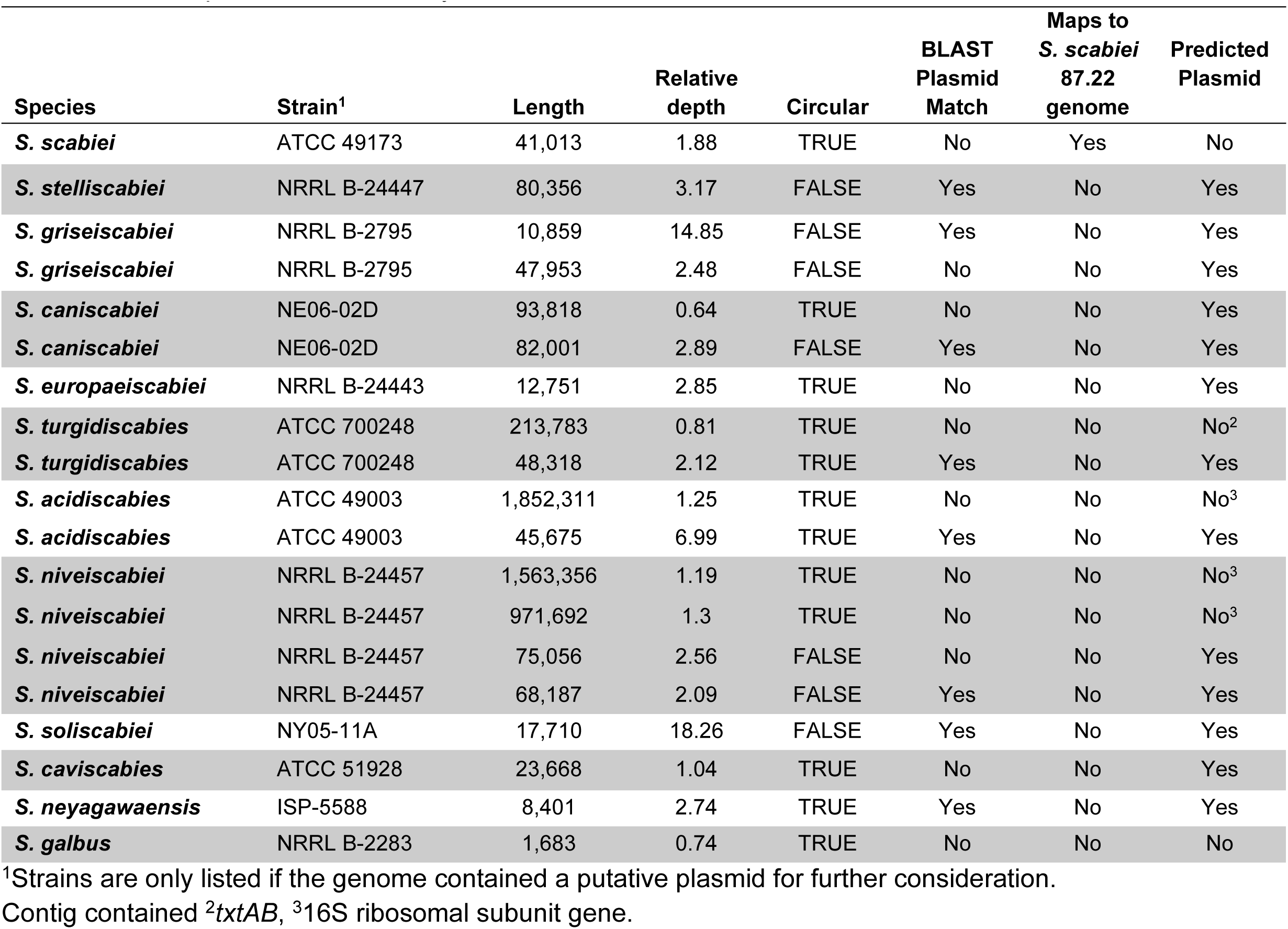
Putative plasmids in assembly FASTA file.

| Species | Strain<sup>1</sup> | Length | Relative depth | Circular | BLAST Plasmid Match | Maps to *S. scabiei* 87.22 genome | Predicted Plasmid |
| --- | --- | --- | --- | --- | --- | --- | --- |
| *S. scabiei* | ATCC 49173 | 41,013 | 1.88 | TRUE | No | Yes | No |
| *S. stelliscabiei* | NRRL B-24447 | 80,356 | 3.17 | FALSE | Yes | No | Yes |
| *S. griseiscabiei* | NRRL B-2795 | 10,859 | 14.85 | FALSE | Yes | No | Yes |
| *S. griseiscabiei* | NRRL B-2795 | 47,953 | 2.48 | FALSE | No | No | Yes |
| *S. caniscabiei* | NE06-02D | 93,818 | 0.64 | TRUE | No | No | Yes |
| *S. caniscabiei* | NE06-02D | 82,001 | 2.89 | FALSE | Yes | No | Yes |
| *S. europaeiscabiei* | NRRL B-24443 | 12,751 | 2.85 | TRUE | No | No | Yes |
| *S. turgidiscabies* | ATCC 700248 | 213,783 | 0.81 | TRUE | No | No | No<sup>2</sup> |
| *S. turgidiscabies* | ATCC 700248 | 48,318 | 2.12 | TRUE | Yes | No | Yes |
| *S. acidiscabies* | ATCC 49003 | 1,852,311 | 1.25 | TRUE | No | No | No<sup>3</sup> |
| *S. acidiscabies* | ATCC 49003 | 45,675 | 6.99 | TRUE | Yes | No | Yes |
| *S. niveiscabiei* | NRRL B-24457 | 1,563,356 | 1.19 | TRUE | No | No | No<sup>3</sup> |
| *S. niveiscabiei* | NRRL B-24457 | 971,692 | 1.3 | TRUE | No | No | No<sup>3</sup> |
| *S. niveiscabiei* | NRRL B-24457 | 75,056 | 2.56 | FALSE | No | No | Yes |
| *S. niveiscabiei* | NRRL B-24457 | 68,187 | 2.09 | FALSE | Yes | No | Yes |
| *S. soliscabiei* | NY05-11A | 17,710 | 18.26 | FALSE | Yes | No | Yes |
| *S. caviscabies* | ATCC 51928 | 23,668 | 1.04 | TRUE | No | No | Yes |
| *S. neyagawaensis* | ISP-5588 | 8,401 | 2.74 | TRUE | Yes | No | Yes |
| *S. galbus* | NRRL B-2283 | 1,683 | 0.74 | TRUE | No | No | No |

<sup>1</sup>Strains are only listed if the genome contained a putative plasmid for further consideration. Contig contained <sup>2</sup>*txtAB*, <sup>3</sup>16S ribosomal subunit gene.

### Characterization of genome synteny among Thaxtomin-positive Streptomyces

To visualize and examine synteny among the linear chromosomes of *Streptomyces* pathogens, we generated pairwise, whole-genome alignment dot plots (**Figure S5**) and structural graphs (**Figure 3**) for assemblies with three or fewer contigs that correspond to chromosomes. Both approaches illustrated lower synteny towards the distal ends of the linear chromosomes, including pairwise comparisons between closely related, clade I.a species and more distantly related pathogenic strains. The pattern of high conservation in the central region and reduced synteny at the distal ends of the chromosomes was also observed for the two closely related pairs: 1.) *S. niveiscabiei* NRRL B-24457 with *S. acidiscabies* ATCC 49003, and 2.) *S. turgidiscabies* ATCC 700248 with *S. echiniscabiei* WI04-05B. Interestingly, there were multiple large translocations, duplications, or inversions between *S. niveiscabiei* NRRL B-24457 and *S. acidiscabies* ATCC 49003, despite their close phylogenetic relationship (**Figure 3**). There was a large inversion and predicted translocation of a central chromosomal region conserved in *S. turgidiscabies* ATCC 700248 and *S. echiniscabiei* WI04-05B relative to all other strains (**Figure 3**). This inversion does not align with contig breaks and is not an artifact of contig concatenation direction or order (**Figure S5**), suggesting that it is an evolutionary conserved, large-scale inversion. Another notable translocation is present in the comparison of *S. griseiscabiei* NRRL B-2795 to the other closely related clade I.a common scab pathogens, in which one of the visible misalignments likely corresponds to a translocation, in comparison to a misalignment associated with a contig break (**Figure S5**). Furthermore, the dot alignments also reveal that failure to resolve single-chromosomal contigs was primarily due to unassembled, very small contigs representing the distal ends of the chromosomes.

**Figure 3.**
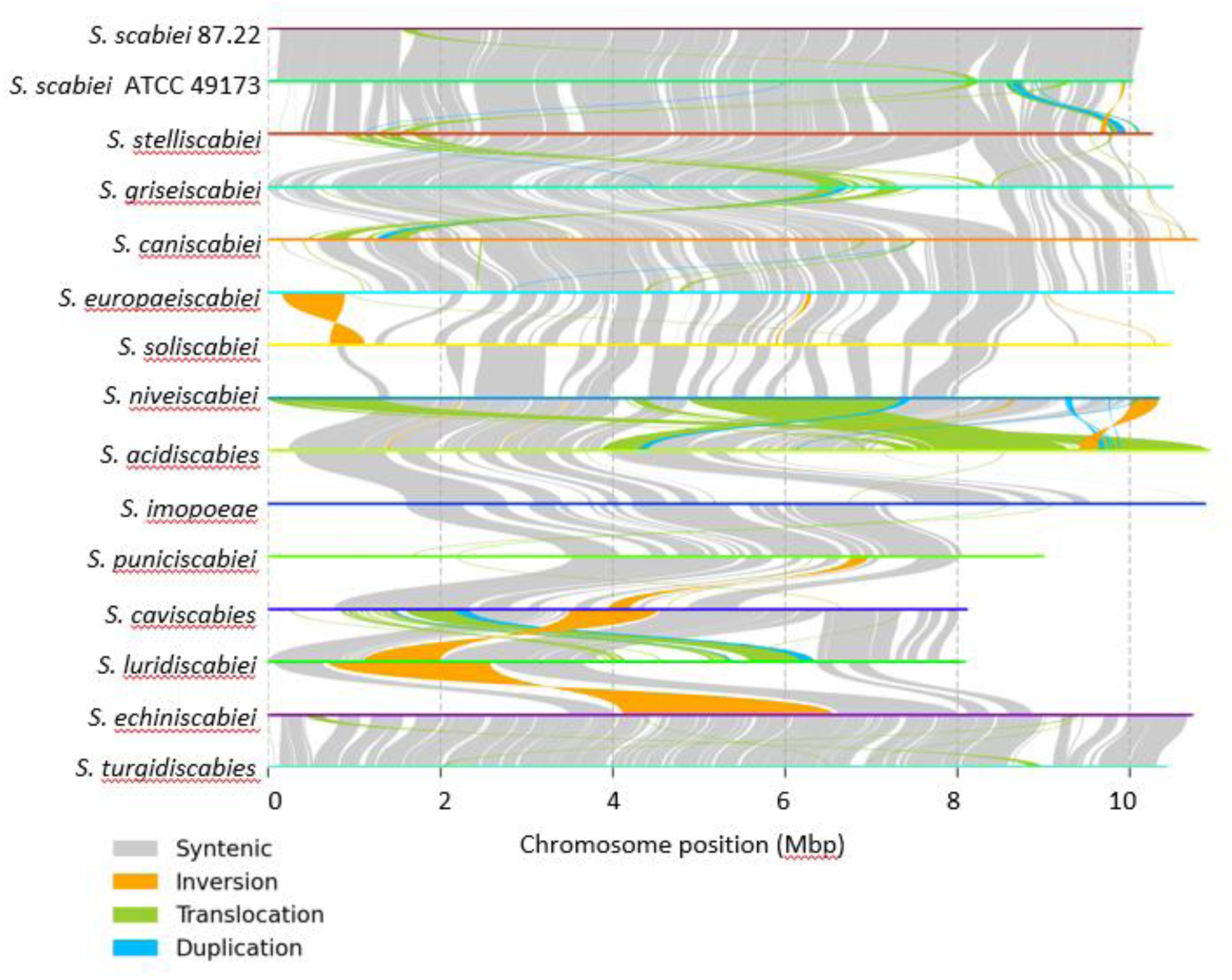
Synteny and genomic rearrangement among common scab- and netted scab-pathogenic strains of *Streptomyces*. Large contigs from genome FASTA files were concatenated into single chromosomes where necessary and ordered pairs of single-chromosome genomes were aligned via nucmer. Alignments were analyzed for synteny and rearrangements via SyRI and plotted via plotsr.

We also investigated the chromosomal position of the Thaxtomin biosynthetic gene-containing TR1 genetic element, a mobilizable element bordered most often by the integration site *aviX1* [50,51]. In most *txtA*-positive genomes, the mobile genetic element was located within 3.5-4 Mb from the end of the chromosome (**Figure S6**). The exceptions were *S. turgidiscabies* ATCC 700248 and *Streptomyces* sp. WI04-05B, in which the Thaxtomin A biosynthetic genes were nearer to the ends of the chromosome. These two strains were previously characterized to possess different TR1 integration sites than the other phytopathogenic strains [50].

## Discussion

Here we report complete or improved, near-complete high-quality genome sequences for 12 pathogenic type-strains of *Streptomyces*, as well as 6 type-strains of non-pathogenic species of sister taxa. The genomes of *Streptomyces* that contain the Thaxtomin A biosynthetic genes and correspond to common scab pathogenicity are the largest of all publicly available complete *Streptomyces* genomes, predicted to have between 9,000 and 10,000 genes (**Figure 1B**). Interestingly, among the non-common scab-causing pathogenic *Streptomyces,* only *S. ipomoeae* strain NRRLB-12321 contains a similarly large gene count. Most *S. ipomoeae* strains contain genes involved in thaxtomin biosynthesis, but these strains do not produce Thaxtomin A, instead produce one or more toxins that correspond to lower disease severity on potato [13].

Pangenome analysis of 153 complete genome assemblies of *Streptomyces* revealed that the genetic diversity of the genus is underpinned by more than 98,000 unique gene families. In general, many genes were only present in one or few genomes, indicating tremendous genetic diversity across the genus (**Figure S3A**). While Thaxtomin A and the presence of the biosynthetic genes are known to be essential for common scab disease [2,6,7,52], additional genes have been acquired that may also be linked to pathogenicity. GWAS and KEGG analyses suggested that phytopathogenicity in common scab-causing species of *Streptomyces* may be associated with the presence of genes involved in functions such as xenobiotics degradation and biosynthesis of secondary metabolites (**Figure 2**). Additionally, genes with predicted functions in simple sugar metabolism and metabolism of aromatic amino acids were shown to be enriched among common scab pathogens. Notably, Thaxtomin is synthesized from two aromatic amino acids: tryptophan and phenylalanine [53], and expression of the Thaxtomin A biosynthetic genes has been shown to be regulated by the sugar cellobiose, as well as aromatic amino acids [6,52,54]. Numerous secondary metabolite biosynthetic genes were enriched among genomes of pathogenic strains, and other specialized metabolites have been associated with virulence [55]. Gene families that have been identified as unique to the pathogenic strains and that play a direct role in virulence still remain unknown.

### Plasmids are not necessary for common scab disease

Unlike other Gram-positive plant pathogens (e.g., *Clavibacter michiganesis*, *Clavibacter capsici,* and *Rhodococcus* spp.), plasmids, while present in some *Streptomyces* strains, do not appear to be an important aspect of pathogenicity for common scab pathogens [17,25,26,56,57]. Our analysis of the newly sequenced genomes suggested that there are no conserved plasmids across the pathogenic strains, and that several phytopathogenic strains contain no putative plasmids (**Table 3**). This lack of association between plasmids and pathogenicity was previously shown for the closely related, soil rot-causing, *Streptomyces ipomoea* pathogens [24]. Generally, the Thaxtomin A biosynthetic genes were found on the largest contig (i.e., chromosome) in each assembly, except for ATCC 700248^T^, for which the Thaxtomin A biosynthetic genes were localized to a 213,783-bp contig. However, this contig was not identified in the PacBio long-read-only assembly for ATCC 700248, suggesting that it is an assembly artifact due to repetitive elements, which can also give rise to greater sequencing depth. Even though super plasmids have been previously identified in *Streptomyces,* all contigs in this analysis larger than 100 kbp were determined unlikely to be plasmids on the basis of containing known housekeeping genes or mapping to the chromosome of the reference strain *S. scabiei* 87.22 (**Table 3**).

The genomes of *Streptomyces* present several challenges to sequencing, such as high G+C content, the relatively large genome size, and the prevalence of repetitive elements [28]. Poorly assembled genomes potentially misconstrue chromosomal structure and the presence or absence of plasmids. While mega-plasmids have been reported in other *Streptomyces* species [58], repetitive elements likely explain the circularization of the two large contigs, observed in *S. acidscabies* ATCC49003 and *S. niveiscabiei* NRRLB-24457, which give no other indications of being true plasmids and contains sequences that encode known essential housekeeping genes. Another challenge to the identification of plasmids from predicted circular contigs was evident in the observation of 10,859- (*S. griseiscabie*i NRRL B-2795) and 17,710-bp (*Streptomyces* sp. NY05-11A) linear contigs with relative assembly depths of 14.85 and 18.26x, respectively (**Table 3, Figure S4**). Neither of these high copy, putative plasmids contained 16s rRNA genes or other housekeeping genes. Furthermore, these two linear contigs produced BLAST+ alignments to known *Streptomyces* plasmids, further supporting their designation as linear plasmids, which have been reported in other *Streptomyces* species [59–61]. Such challenges affirm the utility of generating complete and nearly complete genomes of *Streptomyces*.

### The distal ends *of Streptomyces* linear chromosomes have lower syntenic conservation

The sequenced *Streptomyces* genome assemblies produced an average of 1-3 contigs comprising the linear chromosome (**Figure S4**), as reported for other *Streptomyces* spp. [22,60,62]. Whole chromosome alignments and dot plots of the complete genome assemblies of the newly sequenced genomes demonstrated an overall conservation of chromosome structure among closely related species, but genome synteny is less conserved at the distal ends of the linear chromosomes (**Figure 3**), as previously reported with select strains in the genus [63]. These data also reveal multiple rearrangements and inversions among the genomes. Breaks in the synteny plots occurred most often in flanking regions of the chromosome, matching earlier reports that chromosome rearrangement is highest in the telomeric regions [64].

## Conclusions

These improved genome assemblies provide the necessary groundwork to explore further features associated with common scab pathogenicity and characterize the diversification of phytopathogenic *Streptomyces*. Using only complete or nearly complete genomes facilitated confirmation that the phytopathogenic *Streptomyces* have much larger genomes on average than non-phytopathogenic *Streptomyces* and supported the identification of gene functions and pathways that appear to be enriched specifically in the genomes of the pathogenic strains. These complete or nearly complete genome assemblies of pathogenic *Streptomyces* also enabled comparisons of chromosomal structure across species and supported the finding that plasmids are not a defining feature of *Streptomyces* pathogenicity.

## List of abbreviations

BUSCO: Benchmarking Universal Sing-Copy Orthologs
CDS: coding sequence
MLSA: multi-locus sequence analysis
SRA: sequence read archive
GWAS: genome-wide association study

## Data reporting

All sequence data generated by this project has been deposited under NCBI Bioproject PRJNA1193568, and accession numbers are listed in Table 1.

## Author contributions

BAS acquired and analyzed data and wrote the manuscript. MLF acquired and analyzed data and wrote the manuscript. HPN acquired data. AJW acquired and analyzed data. JHC analyzed data. CRC designed the work, analyzed data, and wrote the manuscript. All authors read and approved the final manuscript.

