## Supplemental figures for "Improved genome assemblies of plant-associated *Streptomyces* spp. as a resource for understanding plant pathogenicity in the genus"

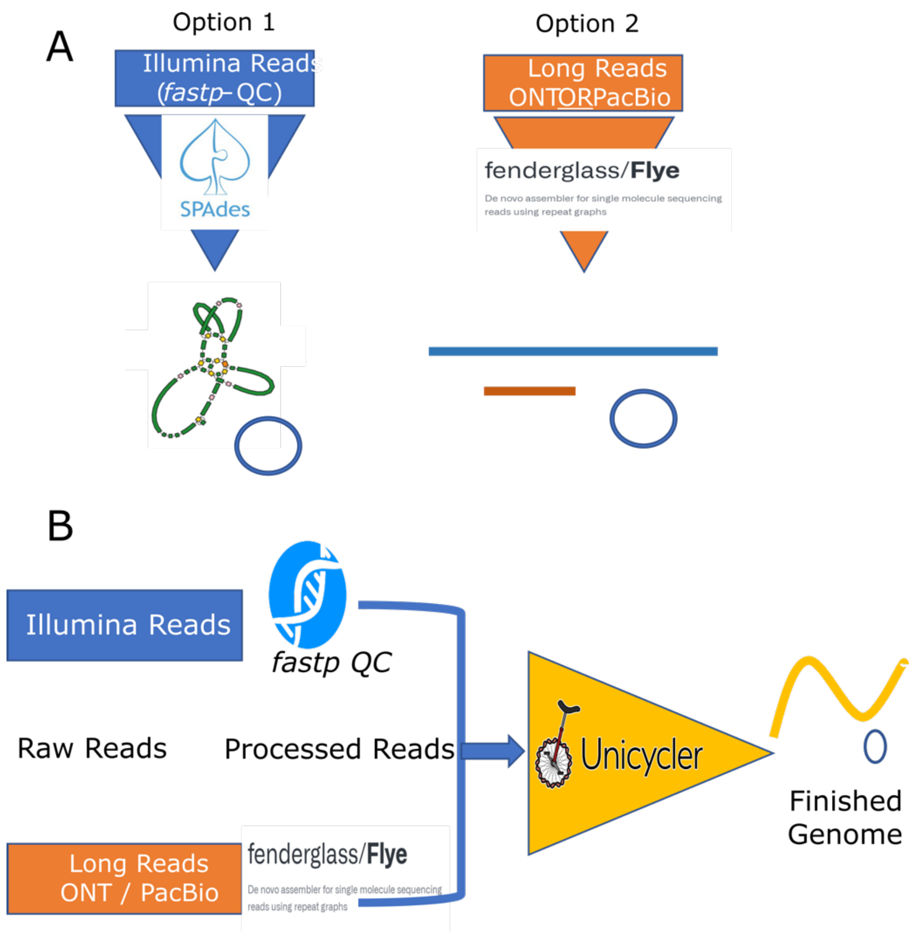


**Figure S1. Illustration of the workflow implemented to generate complete genomes of the phytopathogenic *Streptomyces* type strains. (A)** Methods used for generating genome assemblies from single-source sequencing technology reads. **(B)** Hybrid assembly pipeline used in this work to generate near-complete genomes of *Streptomyces* strains.

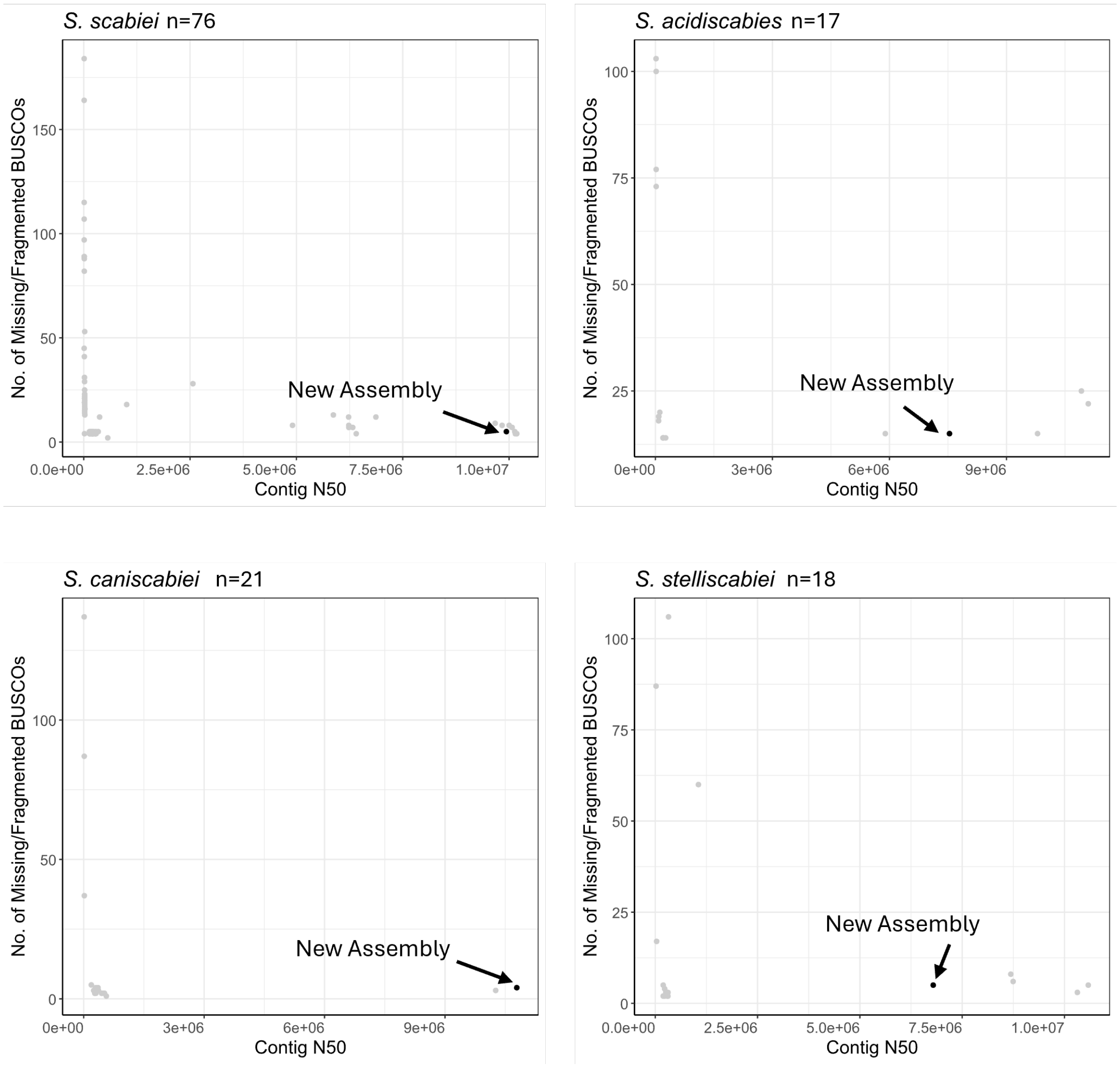


**Figure S2. Contig N50 and missing or fragmented BUSCOs among the four most highly sequenced species groups of phytopathogenic *Streptomyces*.** All available strains for each species group were downloaded from NCBI on May 17, 2024. The new type strain genome assembly in this work of each species is indicated in grey with an arrow.

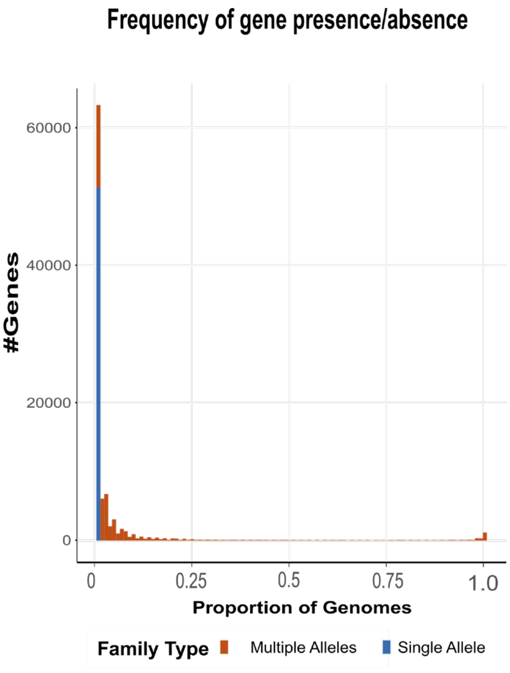

Frequency of gene presence/absence
#Genes
60000
40000
20000
0
0
0.25
0.5
0.75
1.0
Proportion of Genomes
Family Type
Multiple Alleles
Single Allele


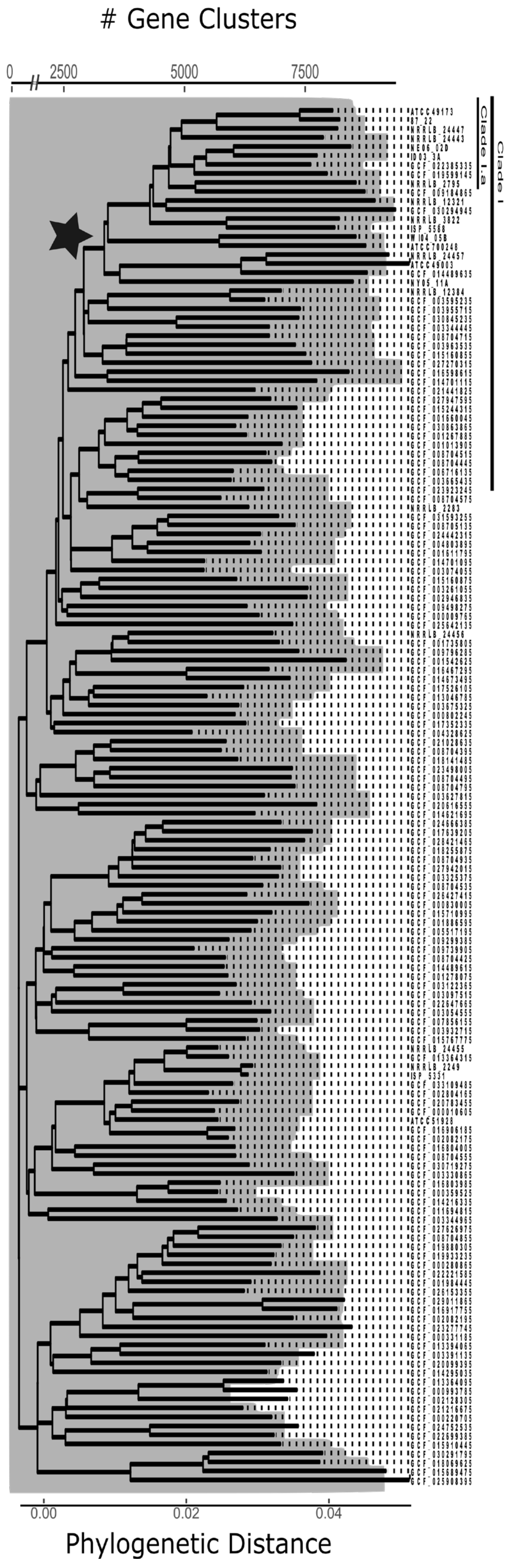

### Gene Clusters
0
2500
5000
7500
Clade I.a
Clade I
0.00
0.02
0.04
Phylogenetic Distance

**Figure S3. Pangenome composition and phylogenetic distribution of gene families in complete and near-complete *Streptomyces* genome assemblies. (A)** Histogram of number of the orthologous gene families. Gene families are considered stable (when they have only a single orthologs at 98% amino acid identity - blue) or diverged (when they had >1 allele for each orthologous gene- red). **(B)** Overlay of the number of identified gene clusters per genome reported from PIRATE for all *Streptomyces* species with complete genome assemblies onto the phylogenetic tree. Pathogenic species are sub-divided into Clade I.a depending on Thaxtomin A PAI. Node where genome enlargement occurs is marked by star.

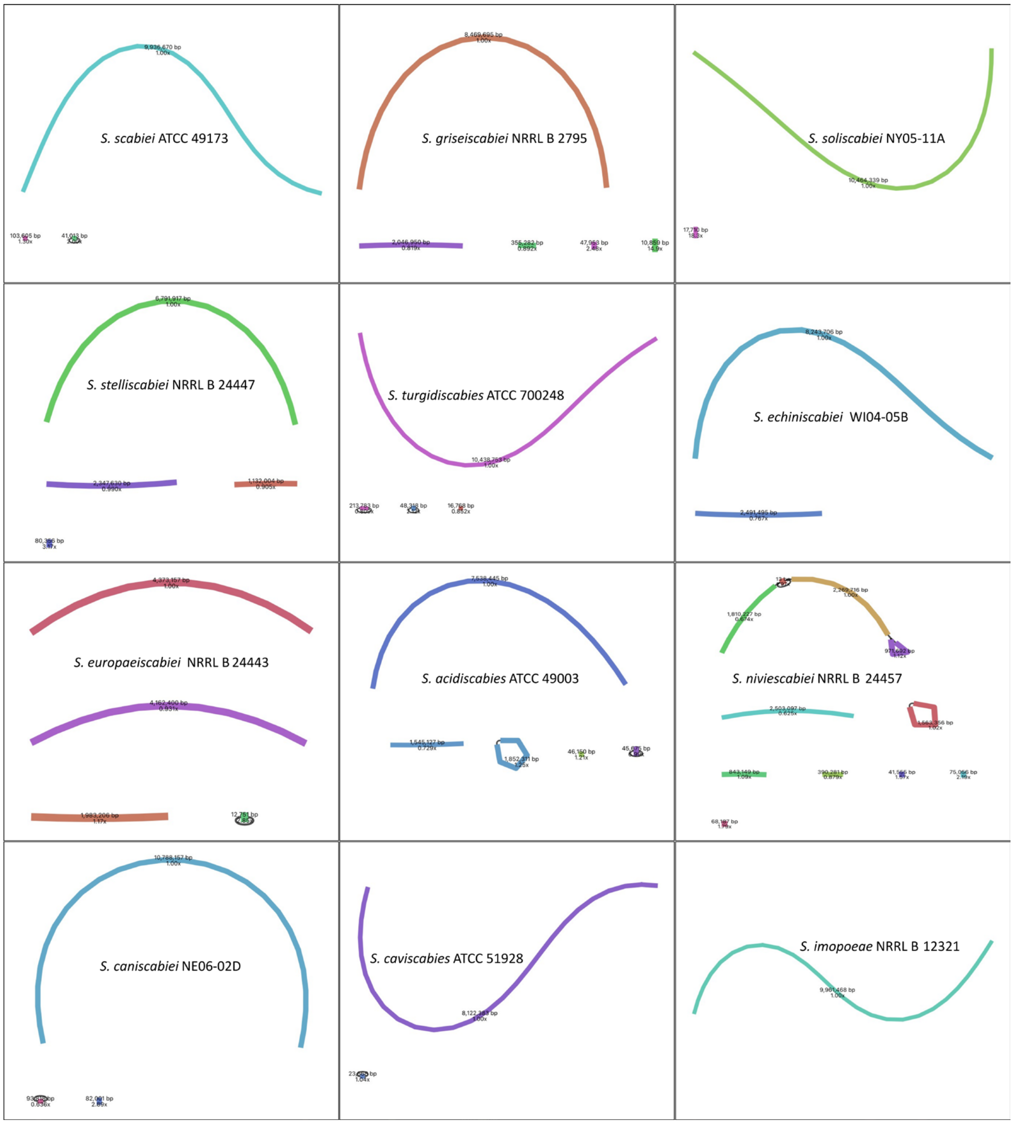


**Figure S4. Bandage graphs of genome assemblies for the pathogenic species reported in this paper.** *Streptomyces* species and strain names are indicated in each block with the number of contigs (lengths). Circular contigs and other putative plasmids are reported in Table 3.

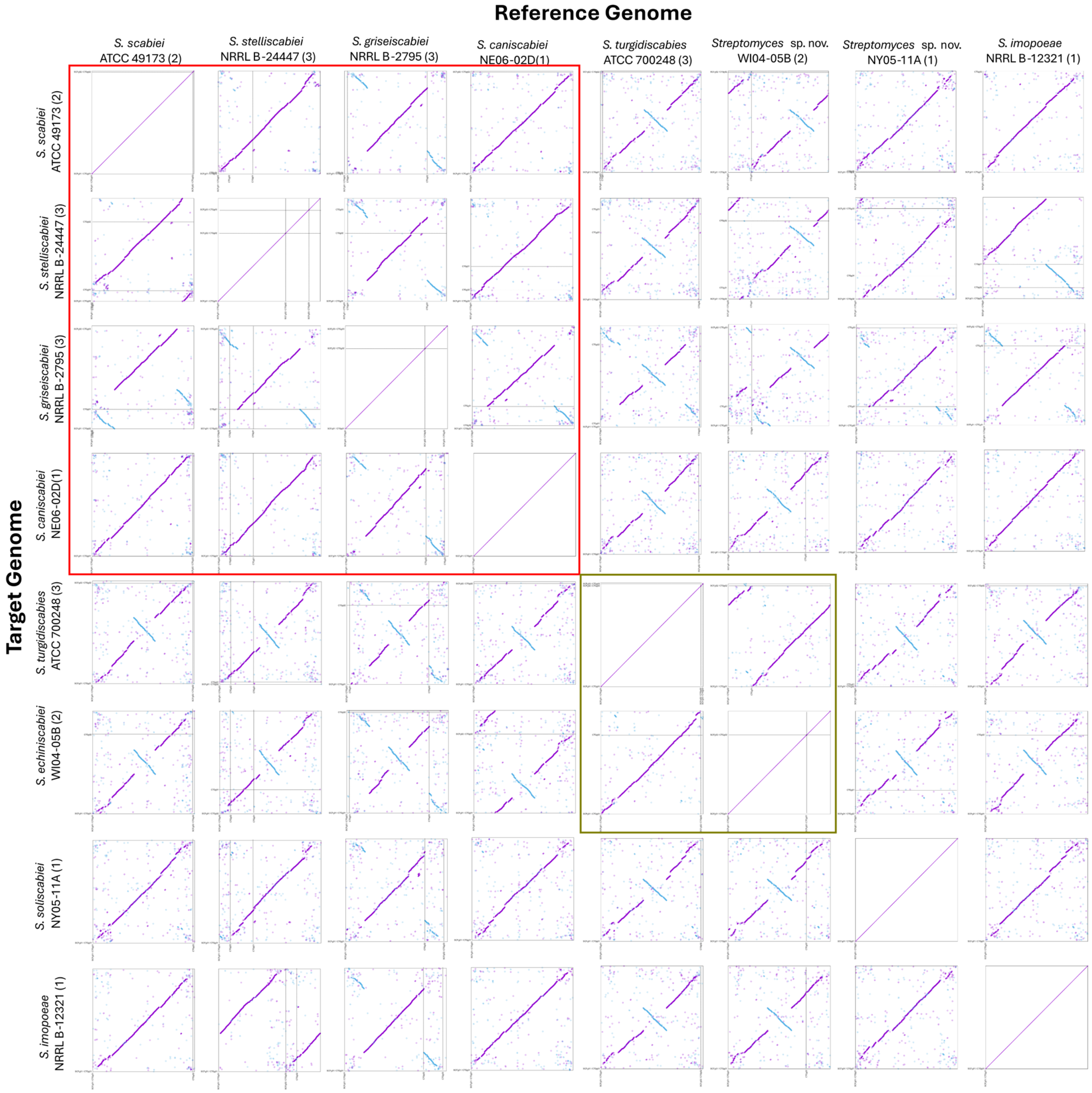


**Figure S5. Dot plots of complete *Streptomyces* genome assemblies.** Genomes were mapped using CSAR (contig scaffolding using algebraic rearrangements) webserver [3]. Dot plots of closely related *Streptomyces* to four closely related clade I.a pathogenic *txtA*-positive strains and four more distantly related common scab pathogens. Contigs boundaries are named on the bottom and left of each panel with dashed lines. Number of contigs is listed in parentheses next to strain names. Sequences of putative plasmids were removed from genome sequences prior to alignment. The closely related clade I.a pathogens are highlighted by a red box. Green box indicates closely related strains *S. echiniscabiei* WI04-05B and *S. turgidiscabies* ATCC700248 that share a large inversion relative to the other genomes.

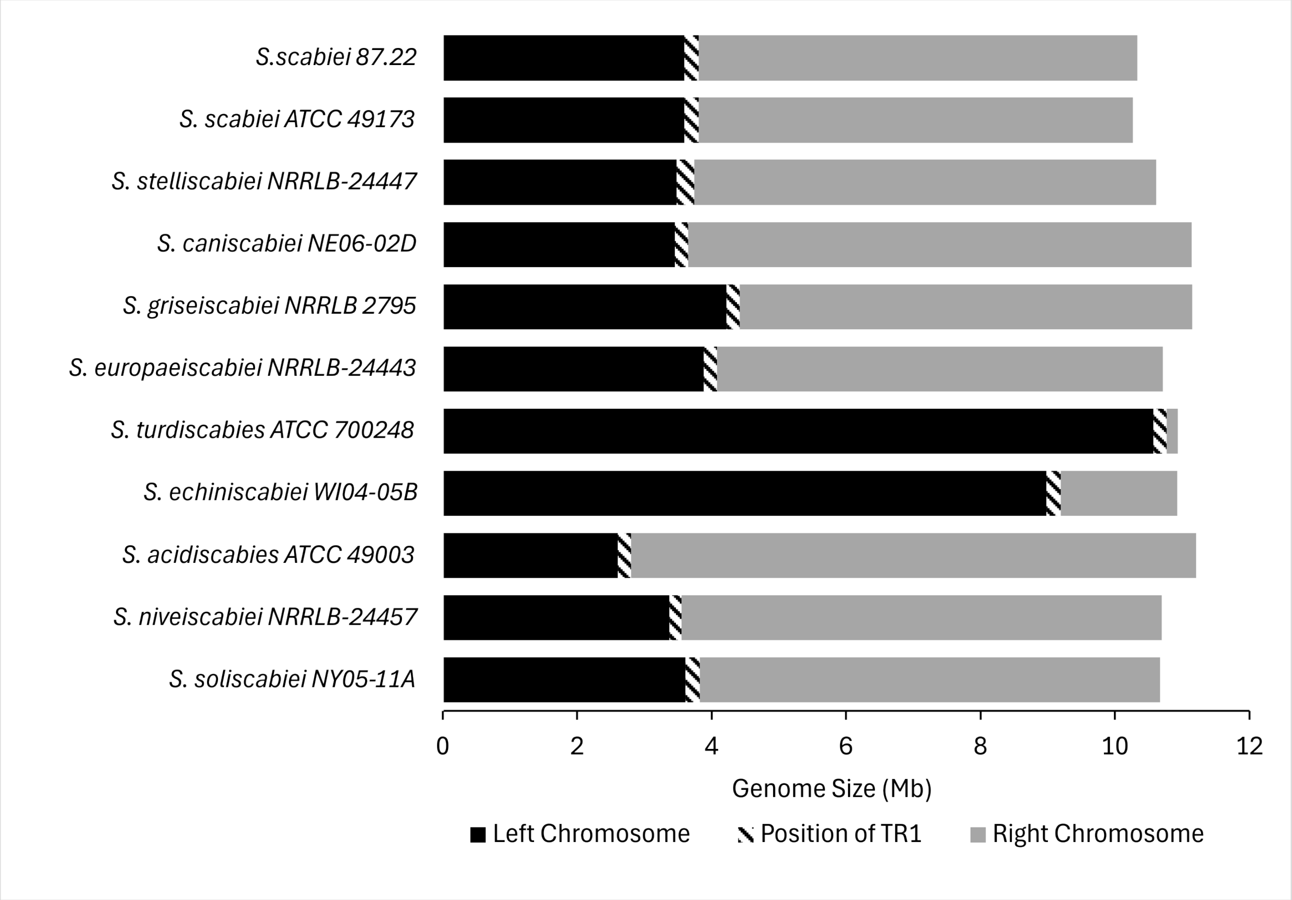


**Figure S6.** Position of the 20-kb TR1 (Thaxtomin A biosynthetic gene cluster) region along the chromosome of the newly finished genome assemblies from 10 *Streptomyces* species. Nearly finished genome assemblies were aligned to the reference genome for *S. scabiei*, strain 87.22, using progressiveMauve [33]. Contigs that were inverted in the target genome assemblies were manually reverse complemented and re-ordered to match the placement along the 87.22 genome assembly.
